# A latent variable model for evaluating mutual exclusivity and co-occurrence between driver mutations in cancer

**DOI:** 10.1101/2024.04.24.590995

**Authors:** Ahmed Shuaibi, Uthsav Chitra, Benjamin J. Raphael

**Author notes:** These authors contributed equally (Ahmed Shuaibi, Uthsav Chitra).

## Abstract

A key challenge in cancer genomics is understanding the functional relationships and dependencies between combinations of somatic mutations that drive cancer development. Such *driver* mutations frequently exhibit patterns of *mutual exclusivity* or *co-occurrence* across tumors, and many methods have been developed to identify such dependency patterns from bulk DNA sequencing data of a cohort of patients. However, while mutual exclusivity and co-occurrence are described as properties of driver mutations, existing methods do not explicitly disentangle functional, driver mutations from neutral, *passenger* mutations. In particular, nearly all existing methods evaluate mutual exclusivity or co-occurrence at the gene level, marking a gene as mutated if any mutation – driver or passenger – is present. Since some genes have a large number of passenger mutations, existing methods either restrict their analyses to a small subset of suspected driver genes – limiting their ability to identify novel dependencies – or make spurious inferences of mutual exclusivity and co-occurrence involving genes with many passenger mutations. We introduce DIALECT, an algorithm to identify dependencies between pairs of *driver* mutations from somatic mutation counts. We derive a latent variable mixture model for drivers and passengers that combines existing probabilistic models of passenger mutation rates with a latent variable describing the unknown status of a mutation as a driver or passenger. We use an expectation maximization (EM) algorithm to estimate the parameters of our model, including the rates of mutually exclusivity and co-occurrence between drivers. We demonstrate that DIALECT more accurately infers mutual exclusivity and co-occurrence between driver mutations compared to existing methods on both simulated mutation data and somatic mutation data from 5 cancer types in The Cancer Genome Atlas (TCGA).

## 1 Introduction

Cancer is an evolutionary process driven by a small number of somatic *driver* mutations against a larger background of random and functionally neutral (or slightly deleterious) *passenger* mutations [28, 80, 49]. Distinguishing driver mutations from passenger mutations and understanding the function of driver mutations is critical for understanding cancer progression and for developing targeted cancer therapies [25] To this end, large-scale sequencing projects such as the International Cancer Genome Consortium (ICGC) [32,81] and The Cancer Genome Atlas (TCGA) [51, 9, 44, 37, 76, 5] have measured somatic mutations in large cohorts of tumor samples, allowing for the systematic analysis of driver mutations across many different cancer types.

Beyond the prioritization of individual driver mutations and genes, another important problem in cancer genomics is understanding the functional relationships and dependencies between *combinations* of driver mutations. For example, it has been empirically observed that certain pairs or sets of driver mutations are *mutually exclusive*, meaning that these driver mutations are observed in the same tumor sample less frequently than expected by chance [78]. A widely held explanation for such observed mutual exclusivity is that driver mutations are grouped into a small number of biological pathways, such that a single driver mutation is sufficient to perturb a pathway in a tumor. Combined with the relatively small number of driver mutations in a single tumor, two driver mutations rarely occur in the same pathway. For example, driver mutations in the *KRAS* and *BRAF* genes – two oncogenes in the Ras/Raf/MAP-kinase signaling pathway – have been observed to be mutually exclusive across large cohorts of colorectal cancer samples [18, 7]. Another explanation for mutual exclusivity is synthetic lethality where a pair of mutations – but not the individual mutations – result in cell death [56, 34]. On the other hand, some pairs or sets of driver mutations are *co-occurring*, meaning that they are observed in the same tumor sample more often than expected, e.g. the *VHL/SETD2/PBRM1* mutations in renal cancer [73]. Co-occurrence between driver mutations is observed to be much rarer than mutual exclusivity [10] and may result from some pathways requiring multiple mutations to be perturbed [72].

Numerous computational methods have been developed over the past decade to identify pairs (or larger sets) of genes with mutually exclusive or co-occurring mutations (reviewed by [63, 70, 53]). Importantly, although dependency relationships such as mutual exclusivity and co-occurrence are often described as properties of individual driver mutations, the typical practice is to analyze these dependencies at the *gene* level, treating all observed nonsynonymous single-nucleotide mutations in a gene identically [52, 72, 41, 43, 15, 10, 42, 68, 36, 16, 35,2,45]. (Some methods also analyze larger alterations such as copy number aberrations (CNAs) or DNA methylation changes [59, 41, 10], but we restrict our attention to single nucleotide somatic mutations, which are the vast majority of somatic mutations analyzed by existing methods.) There are three major reasons why mutual exclusivity and co-occurrence analysis is typically performed at the gene level. First, it is often unknown *a priori* which somatic mutations are driver mutations and which are passenger mutations, and the classification of mutations as drivers or passengers remains an active area of research [63]. Second, beyond a small number of mutational hotspots [74], individual genomic positions are mutated infrequently in the available cohorts of hundreds to thousands of patients. Third, it is computationally intractable to analyze all combinations of somatic mutations in a cohort, as most cancers are estimated to contain 1,000-20,000 somatic mutations [48].

Methods for identifying dependencies between driver mutations at the gene level do not explicitly account for passenger mutations. Instead, existing methods typically aggregate all somatic mutations in a gene – both drivers *and* passengers – into a single mutational event. Most of these methods use *ad hoc* procedures to restrict analysis to a small subset of genes that are predicted to be driver genes. However, requiring such prior knowledge substantially limits the ability of these methods to identify novel sets of mutually exclusive or co-occurring driver mutations. On the other hand, if existing methods are used to analyze larger lists of genes, then these methods will identify many *spurious* dependencies involving non-driver mutations. For example, we show that existing methods often identify mutual exclusivity involving mutations in the genes *TTN* or *MUC16*, two genes which are hypothesized to not carry any driver mutations and instead have large numbers of passenger mutations due to their length (>60,000 base-pairs) and high background mutation rates [40]. This empirical observation suggests that separately modeling driver and passenger mutations is a promising approach for identifying dependencies between drivers.

Separately, there is a large line of work on identifying individual driver genes from somatic mutation data (e.g. [69, 20, 40, 75, 67, 21, 27, 30, 4, 55, 26, 3, 13, 12]). Some of these algorithms implicitly (or explicitly) model the number the number of passenger mutations inside each gene, i.e. a *background mutation rate model*, and they identify individual genes whose number of observed somatic mutations is significantly greater than expected under the background mutation model. Critically, such algorithms do not identify genes like *TTN* or *MUC16* as driver genes, as they derive background mutation models using genomic features correlated with increased passenger mutation rates including gene length, replication timing, and synonymous mutation rate [40]. However, these algorithms only model the distribution of passenger mutations inside individual genes, and have not been used to model the distribution of *driver* mutations inside pairs or larger sets of genes.

We introduce a new algorithm, Driver Interactions and Latent Exclusivity or Co-occurrence in Tumors (DIALECT), to identify pairs of genes with mutually exclusive and co-occurring *driver* mutations. We derive a latent variable model for dependencies between driver mutations in a pair of genes, which combines existing probabilistic models of background mutation rates with latent variables that describe the presence or absence of driver mutations in each gene. Importantly, by incorporating existing background mutation rate models, we identify combinations of driver mutations *de novo*; unlike existing approaches, we do not need ad hoc heuristics to analyze small subsets of previously studied driver genes. We derive an expectation-maximization (EM) algorithm to learn the parameters of our model, which describe the rates of mutual exclusivity and co-occurrence between a pair of driver mutations. We use DIALECT to identify dependencies in simulated data and to identify pairs of genes with mutually exclusive driver mutations in real somatic mutation data across 5 cancer subtypes. We show that DIALECT has improved statistical power and lower false positive rate compared to existing methods.

## 2 Methods

We derive a latent variable model for evaluating mutual exclusivity and co-occurrence between driver mutations in a pair of genes. We assume we are given as input a count matrix ***C*** = [*c*_*ij*_] ∈ ℝ^*N* ×*G*^ indicating the number of non-synonymous somatic mutations in *G* genetic loci (e.g. genes) across *N* tumor samples. We aim to test whether each pair (*j, j* ^′^) of genes has mutually exclusive driver mutations. For ease of notation, we omit the subscripts *j* and focus our exposition on a single pair of genes, where the first gene has somatic mutation counts ***c*** = [*c*_*i*_] ∈ ℝ^*N*^ and the second gene has somatic mutation counts 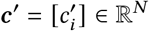.

Let *C*_*i*_ and 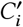 be random variables indicating the number of somatic mutations observed in two genes, respectively, in tumor sample *i* = 1,…, *N*. We assume the somatic mutation count *C*_*i*_ (resp.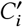) in each sample *i* is equal to the sum of two independent random variables: (1) the number *P*_*i*_ (resp.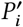) of *passenger* mutations in sample *i*, and (2) an indicator variable *D*_*i*_ ∈ {0, 1} (resp.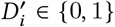) describing the presence or absence of a *driver* mutation in the gene in sample *i*, i.e.

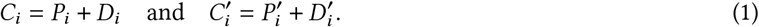

We note that we assume that there is at most one driver mutation in a gene in a given sample, which is a reasonable assumption in many cases^1^.

We aim to estimate the joint distribution 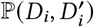 of driver mutations, which describes *dependencies* between driver mutations, i.e. when the random variables *D*_*i*_ and 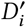 are *not independent*. For example, mutual exclusivity (ME) corresponds to 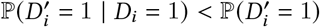 while co-occurrence (CO) corresponds to 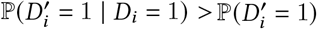. (Note that if *D*_*i*_ and 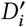 are independent, then 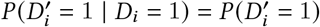.

We emphasize that existing methods do not model the distribution 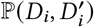 of driver mutations. Instead, these methods first binarize the somatic mutation counts, forming the matrix ***X*** = [*x*_*ij*_] where 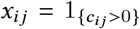, and then analyze the binarized mutation counts ***x*** = [*x*_*i*_] ∈ {0, 1}^*N*^ and 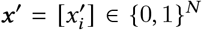 for a pair of genes, respectively (Figure 1A-C). Typically, each binarized counts *x*_*i*_ (resp.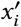) is modeled as a sample of a random variable *X*_*i*_ (resp.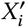), and one aims to test whether the random variables *X*_*i*_ and 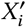 are independent. For example, a classical approach for testing CO and ME is Fisher*’*s exact test, which tests for independence by using a hypergeometric model for the entries of a 2 × 2 contingency table formed from the binarized counts 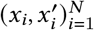.

**Figure 1:**
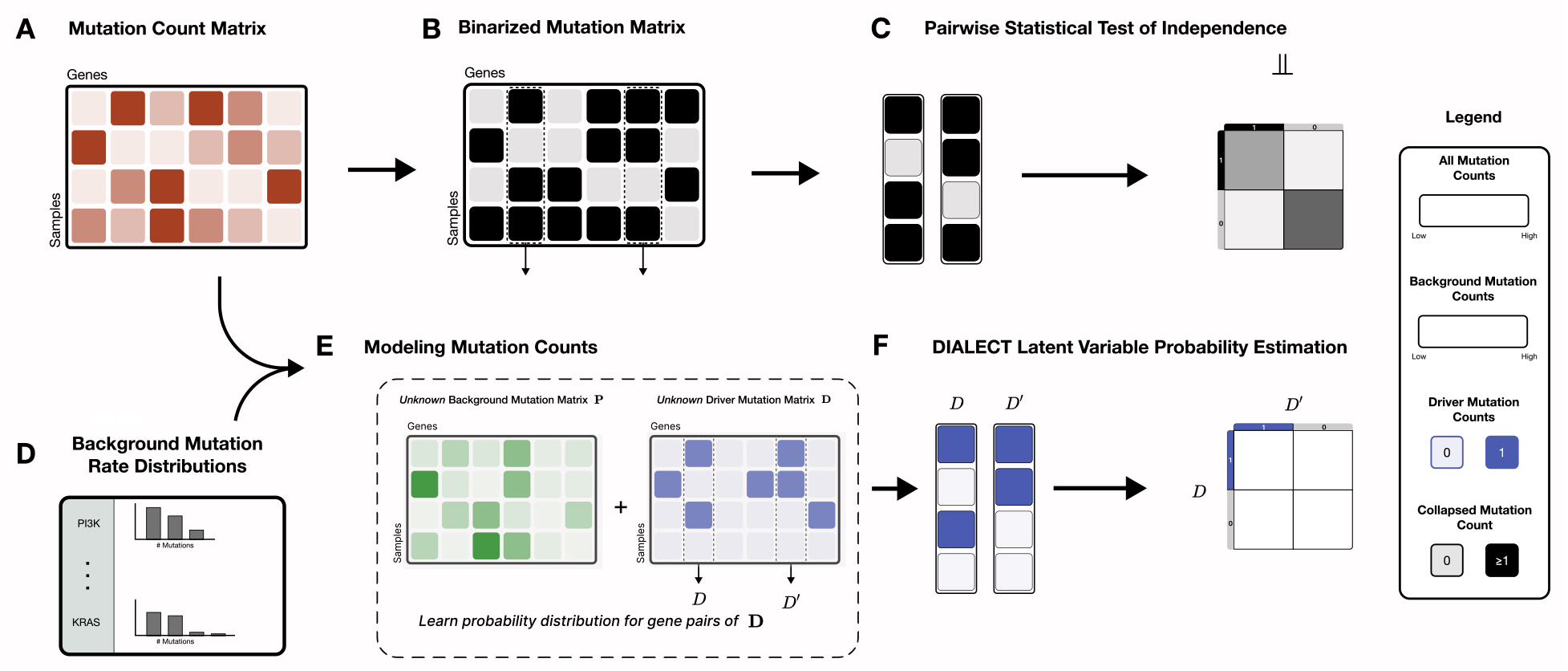
Overview of DIALECT. **(A)** From DNA sequencing data, one obtains a count matrix ***C*** = [*c*_*ij*_] indicating the number of nonsynonymous somatic mutations in genes across tumor samples. **(B)** Existing methods for identifying mutually exclusive driver mutations first create a binarized count matrix 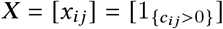 and **(C)** test for independence between pairs of genes. By binarizing the somatic mutation counts, these methods conflate driver mutations versus random, passenger mutations. **(D)** Separately, several algorithms estimate background mutation rate distributions, or the distribution of the number of passenger mutations inside a gene, in order to identify individual driver genes. **(E)** DIALECT explicitly models the distribution of somatic mutation counts *C*_*i*_ = *P*_*i*_ + *D*_*i*_ and 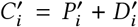 for two genes as a sum of passenger mutations 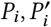, respectively, and latent variables 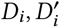, respectively, indicating the presence or absence of driver mutations. DIALECT incorporates background mutation rate distributions ℙ (*P*_*i*_) learned by prior approaches. **(F)** DIALECT learns the parameters τ = (τ_00_, τ_01_, τ_10_, τ_11_) of the driver mutation distribution 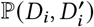 which describes dependencies between drivers including mutual exclusivity and co-occurrence.

The key challenge in estimating the distribution 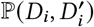 of driver mutations is that we only observe the total number 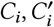 of somatic mutations in a sample and *not* the number 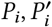 of passenger mutations (or equivalently the value of *D*_*i*_,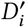). Although the number *P*_*i*_ of passenger mutations is unknown, many methods have been developed to predict driver genes [69, 20, 40, 75, 67, 21, 27, 30, 4, 55, 26, 3, 13, 12] and some of these implicitly (or explicitly) estimate the *distribution* ℙ (*P*_*i*_) of the number *P*_*i*_ of passenger mutations – sometimes called a *background mutation rate* (BMR) distribution (Figure 1D). Note that distributions ℙ (*P*_*i*_) may differ across samples *i* = 1,…, *N* for a variety of reasons, e.g. some tumor samples being hypermutators [65]. In the next section, we show how to use the BMR distributions ℙ (*P*_*i*_) to estimate the distribution of driver mutations.

### 2.1 Driver distribution for a single locus

We start by studying the simple problem of estimating the driver mutation distribution ℙ (*D*_*i*_) in a *single* genetic locus. We will then demonstrate that our approach readily extends to learning the distribution of driver mutations in a pair (or any larger combination) of genetic loci.

We make the simplifying assumption that the driver mutation random variables *D*_*i*_ are *independent and identically distributed* (i.i.d.) across all tumor samples *i* = 1,…, *N*, i.e. the probability of a locus having a driver mutation does not depend on the specific tumor sample. This assumption is motivated by many standard models of tumor growth, where the probability of a cell receiving a driver mutation does not depend on which other mutations are present in the cell [8,23]. The assumption that a particular driver mutation is identically distributed across tumor samples may not always hold, but we demonstrate below that this assumption allows for tractable estimation of the distribution *P* (*D*_*i*_) of driver mutations and works well in practice. Under this assumption, the driver mutations *D*_*i*_ are each independently distributed according to a Bernoulli distribution Bern(*π*) with a shared parameter *π*, representing the *driver mutation rate* across all samples *i* = 1,…, *N*.

Then, the distribution ℙ (*C*_*i*_) of somatic mutation count *C*_*i*_ in sample *i* is given by

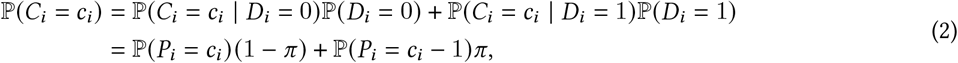

where we use that passenger mutations *P*_*i*_ and driver mutations *D*_*i*_ are independent in the second equation. We set ℙ (*P*_*i*_ = −1) = 0 for notational simplicity, so that the probability of zero somatic mutations in a loci is given by ℙ (*C*_*i*_ = 0) = ℙ (*P*_*i*_ = 0)(1 − *π*). Thus, the log-likelihood ℓ_*C*_ (*π*) = log ℙ (*C*_1_,…, *C*_*N*_; *π*) of the observed somatic mutation counts ***c*** for a gene is given by

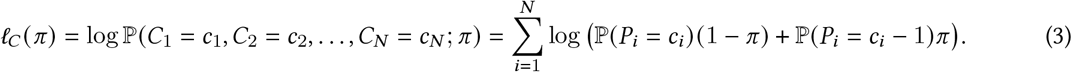

Given observed mutation counts ***c*** and BMR distributions ℙ (*P*_1_),…, ℙ (*P*_*N*_), we compute the driver mutation rate *π* that maximizes the log-likelihood ℓ_*C*_ (*π*) of the observed data:

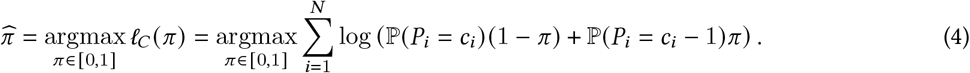

The maximum likelihood problem (4) is challenging to solve exactly as it is often a *non-convex* optimization problem, depending on the form of the background distributions ℙ (*P*_*i*_). We solve this optimization problem by making the observation that the mutation count distribution (2) may be viewed as a *latent variable model*, where the unobserved, binary driver mutations *D*_*i*_ are the *latent variables* and the somatic mutation counts *C*_*i*_ are distributed according to a mixture of two distributions, ℙ (*P*_*i*_) and ℙ (*P*_*i*_ − 1).

The standard approach for computing an MLE for a latent variable model is the *expectation maximization (EM)* algorithm [6]. Thus, we solve (4) using the EM algorithm, whose steps we describe below.

#### E-step

Given an estimated driver mutation rate *π* ^(*t*)^ at iteration *t*, we compute the *responsibility* 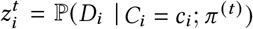, i.e. the probability of the latent variable *D*_*i*_ = 1 being equal to 1 conditioned on the observed mutation count *C*_*i*_, for each sample *i* = 1,…, *N* as

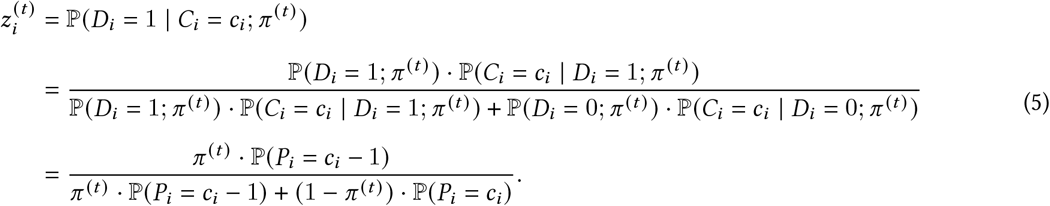

#### M-step

Given the responsibility 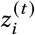 for each sample *i*, we estimate the driver mutation rate *π* ^(*t*+1)^ for iteration *t* + 1 as

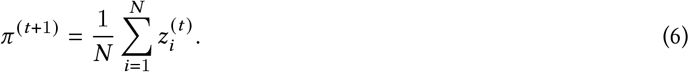

### 2.2 Driver distribution for a pair of loci

We next extend the approach presented above to estimate the distribution 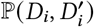 of a *pair* of driver mutations. We start by observing that the driver mutations 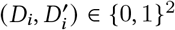 are distributed according to a *bivariate* Bernoulli distribution. A bivariate Bernoulli distribution is specified by four parameters [17]:

1. the probability 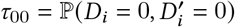 that neither locus has a driver mutation;
2. the probability 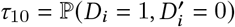 that first locus has a driver mutation;
3. the probability 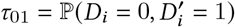 that the second locus has a driver mutation; and
4. the probability 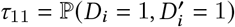 that both loci have driver mutations,

where one of the parameters is redundant since τ_00_ + τ_10_ + τ_01_ + τ_11_ = 1. We note that the bivariate Bernoulli distribution 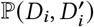 is equivalent to a *categorical* distribution on binary strings 00, 01, 10, 11 with corresponding probabilities τ_00_, τ_01_, τ_10_, τ_11_.

The parameters τ = (τ_00_, τ_01_, τ_10_, τ_11_) of the bivariate Bernoulli distribution 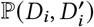 describe whether there is a *statistical interaction* [71] between the driver mutation *D*_*i*_ in the first locus and the driver mutation 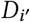 in the second locus. If τ_11_τ_00_ *<* τ_01_τ_10_, then the driver mutations are more likely to be mutually exclusive across samples than not (i.e. a *negative* interaction) while if τ_11_τ_00_ *>* τ_01_τ_10_, then the driver mutations are more likely to co-occur across samples than not (i.e. a *positive* interaction). Driver mutations *D*_*i*_ and 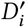 are independent (i.e. no interaction) if and only if τ_11_τ_00_ = τ_01_τ_10_.

More concisely, the interaction between driver mutations is quantified by the *log-odds ratio* 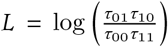, which has previously been previously used to measure ME and CO for binarized mutations [38, 60, 14, 58]. The sign sgn(ℓ) of the log-odds ratio ℓ determines the type of interaction: a positive log-odds ratio *L >* 0 describes ME between the driver mutations 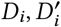 while a negative log-odds ratio *L <* 0 describes CO.

Following a similar derivation as in the previous section, the distribution 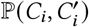 of mutation counts is given by

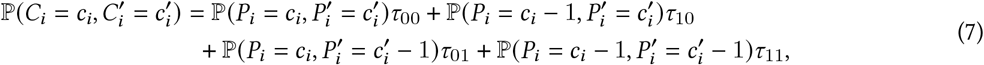

and the log-likelihood 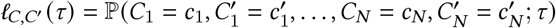 is equal to

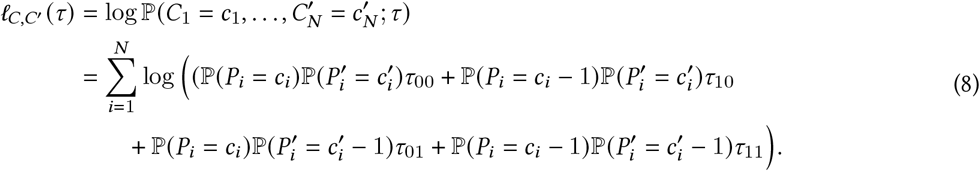

Given observed mutation counts ***c, c***^′^ for a pair of genes and passenger mutation distributions 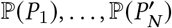 across *N* tumor samples, we compute the parameters τ_00_, τ_01_, τ_10_, τ_11_ of the driver mutation distribution that maximize the log-likelihood of the observed data:

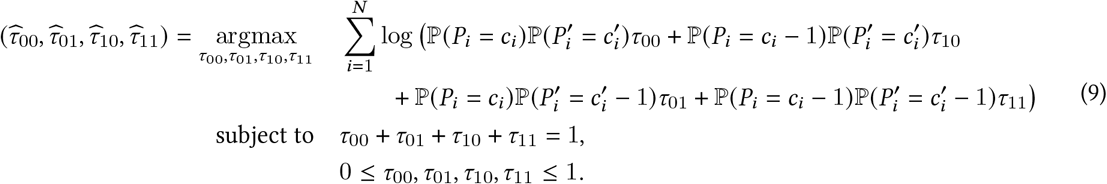

The maximum likelihood problem (9) is difficult to solve as, for many background distributions ℙ (*P*_*i*_), it a non-convex optimization problem over a three-dimensional simplex. Thus, similar to the previous section, we solve (9) using the EM algorithm, whose steps we briefly describe below.

#### E-step

Given the estimated driver mutation probabilities 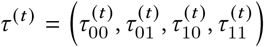 at iteration *t*, we compute the responsibility 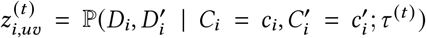 for each driver mutation probability 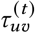 and sample *i* = 1,…, *N* as

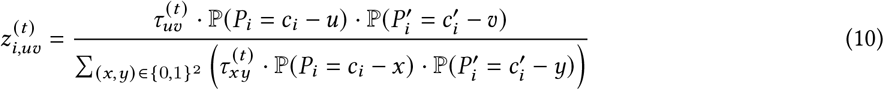

#### M-step

Given the estimated responsibilities 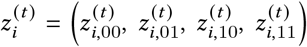 at iteration *t*, we compute the estimated driver mutation probabilities 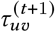 at iteration *t* + 1 as

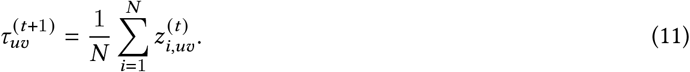

### 2.3 Testing for statistical significance

We test the null hypothesis *H*_0_ that the driver mutations *D*_*i*_, 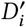 are independent against the alternative hypothesis *H*_1_ that the driver mutations *D*_*i*_, 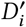 are not independent. We perform this test using the likelihood ratio test (LRT), whose test statistic is equal to the following scalar multiple of the difference between the log-likelihoods under the null hypothesis *H*_0_ and alternative hypothesis *H*_1_:

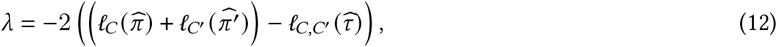

where 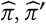 are the estimated driver mutation rates assuming that driver mutations are independent, which are computed by solving (4), and 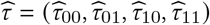 are the estimated parameters of the driver mutation distribution 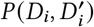 computed by solving (9). We compute a *p*-value assuming that the LRT statistic λ follows a χ^2^-distribution with one degree of freedom, which holds asymptotically by Wilks*’* theorem [77]. We say a pair of genes has ME or CO driver mutations if the *p*-value is less than a threshold *ϵ*.

### 2.4 DIALECT

We implement the EM algorithm for the latent variable model described above in an algorithm called Driver Interactions and Latent Exclusivity or Co-occurrence in Tumors (DIALECT, Figure 1). Given a mutation count matrix C (Figure 1A) and estimated BMR distributions ℙ (*P*_*i*_), 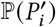 for each gene (Figure 1D), DIALECT estimates the pairwise driver mutation parameters 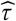 by solving (9) for each pair of genes, and estimates the individual driver mutation rates *π* by solving (4) for each individual gene (Figure 1E-F). DIALECT identifies mutually exclusive (resp. co-occurring) pairs as those with *p*-value less than a threshold *ϵ* (see previous section) and with a positive log-odds ratio 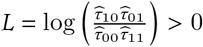 (resp. negative log-odds ratio *L <* 0). We emphasize that the BMR distributions ℙ (*P*_*i*_) used by DIALECT may be estimated using one of several methods, e.g. [40, 75, 67].

## 3 Results

### 3.1 Simulations

We evaluated the ability of DIALECT to identify dependencies between mutations, including mutual exclusivity and co-occurrence, in simulated somatic mutation data.

#### Data

We simulated somatic mutation counts 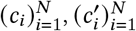 for a pair of genes with lengths *l* and *l* ^′^, respectively, in nucleotides following equation (1). The passenger mutation count *P*_*i*_ (resp.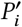) in sample *i* is drawn from a binomial distribution Binom(*l, μ*) (resp. Binom(*l*^′^, *μ*^′^)) where *μ* (resp, *μ*^′^) is a per-nucleotide mutation rate. Such binomial distributions are often used in background mutation rate (BMR) models [40]. We drew each driver mutation 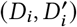 from a bivariate Bernoulli distribution with parameters τ = (τ_00_, τ_01_, τ_10_, τ_11_), where we choose the parameters τ to describe either mutual exclusivity or co-occurrence of driver mutations.

#### Mutual exclusivity

We first assessed DIALECT in identifying *mutually exclusive* driver mutations. We compared DIALECT with two approaches for identifying mutual exclusivity from binarized mutations: Fisher*’*s exact test [22], a classical statistical test of independence; and MEGSA [31], a recent method for identifying mutually exclusive driver mutations.

We simulate somatic mutation counts 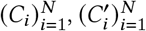 across *N* = 1000 samples with the following parameter choices. The driver mutation distribution 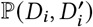 has parameters τ_11_ = 0, i.e. no co-occurrence between drivers, and τ_01_ = τ_10_ = τ, where τ represents the rate of mutual exclusivity between driver mutations. To specify the passenger count distributions, we use gene lengths *l* = *l* ^′^ = 10000 and we use nucleotide mutation rate *μ* = 10^−6^ for the first gene, which was chosen so that the probability ℙ (*P*_*i*_ *>* 0) ≈ 0.01 of this gene having more than one passenger mutation matches the median probability ℙ (*P*_*i*_ *>* 0) across all genes in real data. In order to model how power varies with the presence of passenger mutations, we vary the nucleotide mutation rate *μ*^′^ of the second gene such that that the BMR probability 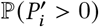, or the probability of the second gene having more than one passenger mutation, varies between 0.01 and 0.10. We assume there are no hypermutated samples, i.e. samples *i* with mutation factor *s*_*i*_ *>* 1.

We run DIALECT with the true BMR distributions ℙ (*P*_*i*_), 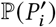 for each sample *i* = 1,…, *N*. Since the power and specificity improves with an increasing number *N* of samples, we choose the *p*-value threshold *ϵ* based on the number *N* of samples: if *N* ≥ 1000 then we set the *p*-value threshold to be *ϵ* = 0.05, while if *N <* 1000 then we set the *p*-value threshold to *ϵ* = 0.001. For Fisher*’*s exact test, a gene pair was identified as mutually exclusive if the resulting *p*-value was less than 0.05. For MEGSA, a gene pair is identified as mutually exclusive if the MEGSA *p*-value, i.e. the MEGSA LRT statistic under the χ^2^-distribution, is less than 0.10.

We observe (Figure 2A) that DIALECT has greater power compared to Fisher*’*s exact test and MEGSA across a range of driver mutual exclusivity rates τ and BMR probabilities 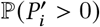. In particular, DIALECT has substantially larger power than Fisher*’*s exact test and MEGSA when the gene pairs have small rates τ of mutually exclusivity (τ ≤ 0.05) and there are a small number of passenger mutations 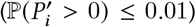— parameters which describe many pairs of driver genes in real data. For these parameter choices, we also performed a *power analysis* and assessed the number of samples needed to achieve a given statistical power. We found (Figure 2B) that *N* > 1000 samples are needed for DIALECT to achieve power > 0.75, while *N >* 2500 samples are needed for Fisher*’*s exact test and MEGSA to achieve the same power. We emphasize that most large cohort studies only measure *N* = 100 − 1000 samples, meaning that DIALECT, as well as existing approaches like Fisher*’*s exact test, may not have sufficient power to detect gene pairs with small rates τ of mutual exclusivity. Nevertheless, our simulations demonstrate that for sufficiently large cohort sizes, DIALECT more accurately identifies pairs of mutually exclusive driver mutations compared to standard approaches.

**Figure 2:**
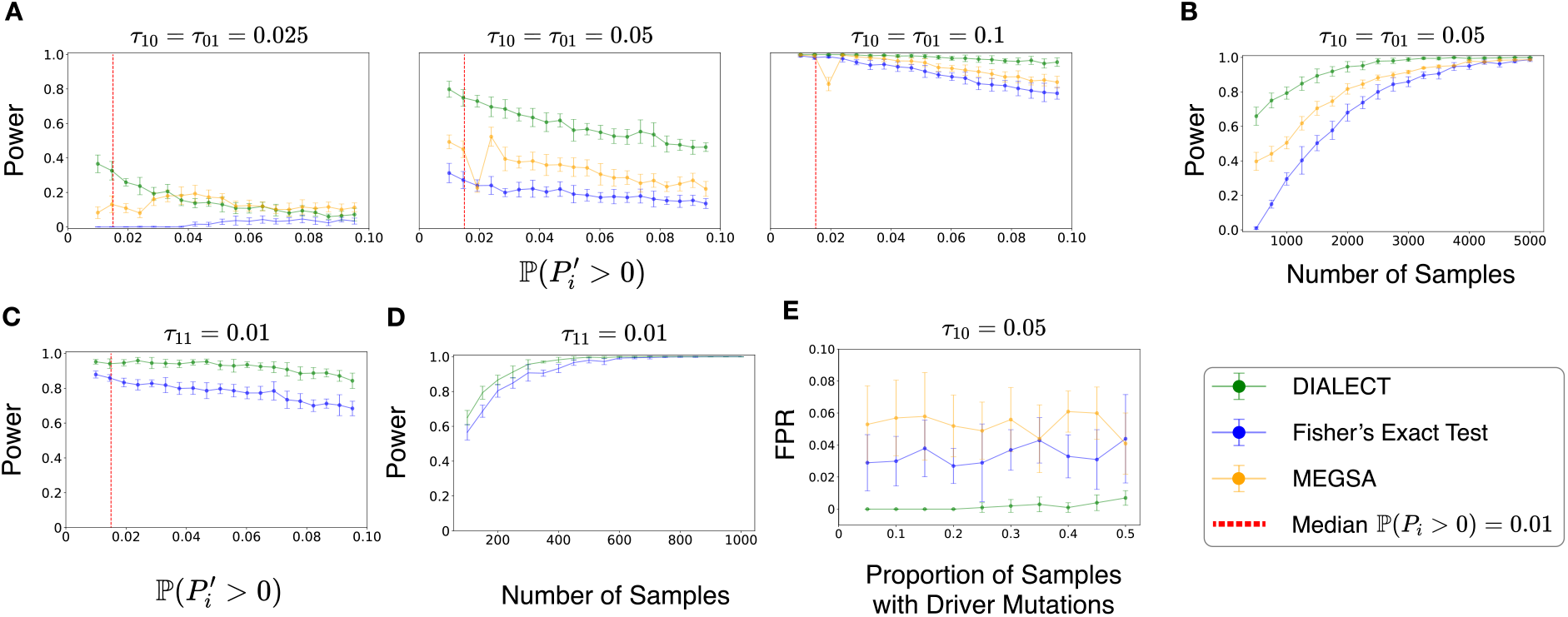
Statistical power and false positive rate for detecting dependencies between driver mutations in simulated data. **(A)** Power (sensitivity) of DIALECT, Fisher*’*s exact test, and MEGSA for identifying mutually exclusive driver mutations from *N* = 1000 tumor samples, for different choices of the rate τ of mutual exclusivity of driver mutations and different probabilities 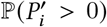 of a gene having passenger mutations. Dashed red line indicates median estimated passenger mutation probability across all genes. **(B)** Power of DIALECT, Fisher*’*s exact test, and MEGSA versus number *N* of samples, which we vary from 100 to 5000, in detecting mutually exclusive driver mutations. **(C)** Power (sensitivity) of DIALECT and Fisher*’*s exact test for identifying co-occurring driver mutations with co-occurrence rate τ_11_ = 0.01 from *N* = 300 tumor samples, for different probability 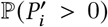 of having passenger mutations. **(D)** Power of DIALECT and Fisher*’*s exact test versus number *N* of samples in detecting co-occurring driver mutations. **(E)** False positive rate versus percentage of samples with driver mutations for τ_10_ = 0.05 across *N* = 1000 samples.

#### Co-occurrence

We next evaluated DIALECT in identifying *co-occurring* driver mutations. We compared DIALECT with Fisher*’*s exact test [22] which tests for co-occurrence in binarized mutations between a pair of genes. We do not compare to MEGSA as it only identifies genes with mutually exclusive mutations. We simulated somatic mutation counts 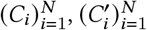 for *N* = 300 tumor samples where (1) the passenger mutation count distributions 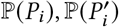 are distributed as previously described and (2) the driver mutation distribution 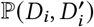 has parameters τ_11_ = 0.01 and τ_01_ = τ_10_ = 0.

We observe that DIALECT has greater power compared to Fisher*’*s exact test across a range of BMR probabilities 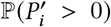 (Figure 2C) and number *N* of samples (Figure 2D). We emphasize that a much smaller number *N* of samples are needed to achieve a power of 1 for identifying co-occurring mutations (*N* ≈ 600, Figure 2D) compared to identifying mutually exclusive mutations (*N* ≈ 5000, Figure 2B), reflecting that co-occurrence is easier to detect than mutual exclusivity. This analysis demonstrates that for small cohort sizes, DIALECT more accurately identifies co-occurring driver mutations than existing approaches.

#### False positive rate

We assessed the false positive rate (FPR, i.e. 1−specificity) of DIALECT and other methods by simulating somatic mutations for a driver gene (i.e. a gene with driver mutations, i.e. *D*_*i*_ = 1 for some samples *i*) and a passenger gene with no driver mutations (i.e.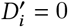) and a large number *P*_*i*_ of passenger mutations. Following the simulation set-up described previously, we set the passenger mutation distribution parameters as *l* = 10000, *μ* = 10^−6^ for the driver gene and *l* ^′^ = 100000 and *μ*^′^ = 10^−5^ for the passenger mutation. The distribution *P* (*D*_*i*_, *D*^′^) of driver mutations has parameters τ_11_ = τ_01_ = 0, and τ_10_ = *π*, where *π* represents the driver mutation rate for the driver gene. Furthermore, in this simulation we assume driver mutations are not identically distributed across samples; instead, we draw driver mutations *D*_*i*_, 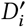 for a *ρ* fraction of all *N* samples selected uniformly at random, where we vary *ρ* between 0.05 and 0.5, and set 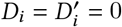 for the other (1 − *ρ*)*N* samples.

We find (Figure 2E) that DIALECT consistently exhibits lower FPR (i.e. higher specificity) than the existing methods across different proportions *ρ* of samples with driver mutations. In particular, DIALECT achieves FPR close to zero when *ρ <* 0.4, which is larger than the mutation rate of nearly all driver genes, while Fisher*’*s exact test and MEGSA have FPR above 0.02. We emphasize that even relatively small FPRs result in the inference of many spurious dependencies in real data analyses. For example, using an algorithm with FPR = 0.01 – which is lower than the FPRs of Fisher*’*s exact test and MEGSA but larger than DIALECT*’*s FPR – to identify dependencies between all pairs of *G* = 100 genes will result in 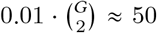 spurious dependencies. We also emphasize that these results show that DIALECT is robust to model mis-specification, since DIALECT assumes driver mutations are identically distributed across tumor samples while our simulated driver mutations are not identically distributed. Such behavior is hypothesized to occur in some cancer types; for example, [70] observed that certain driver mutations are more likely to occur in colorectal cancer subtypes with lower overall mutation loads.

### 3.2 Analysis of mutations in TCGA

We next evaluated DIALECT using somatic mutation data from The Cancer Genome Atlas (TCGA) [76]. We used DIALECT to identify mutual exclusivity, as mutual exclusivity between driver mutations is observed more often than co-occurrence [10, 43]. We compared DIALECT to two state-of-the-art statistical tests for identifying mutual exclusivity: Fisher*’*s exact test [22] and DISCOVER [10]. Fisher*’*s exact test implicitly assumes that each sample is identically distributed, while DISCOVER performs a statistical test where genes have different, sample-specific mutation rates (the DISCOVER test is also asymptotically equivalent to the test used by [42]). However, both Fisher*’*s exact test and DISCOVER use *binarized* mutations as input, and thus do not distinguish between driver mutations and passenger mutations. Since DIALECT analyzes missense mutations and nonsense mutations in a gene separately (since these mutation types often have different background mutation rates), we additionally ran DISCOVER with somatic counts separated into gene events including only nonsynonymous missense mutations (indicated by *GENE_M*) and only nonsense mutations (indicated by *GENE_N*). We denote these results using DISCOVER*. For DISCOVER and DISCOVER* (resp. Fisher*’*s exact test), a gene pair was identified as mutually exclusive if the resulting *q*-value (resp. *p*-value) was less than 0.05.

#### Data

We analyzed non-synonymous mutations from tumor samples in 5 different cancer types from TCGA. Each cancer type contains 100-1000 tumor samples. We obtained the somatic mutation data in Mutation Annotation Format (MAF) from the TCGA PanCancer project, available through cBioPortal [24]. We separately analyzed missense and nonsense mutations, appending gene names with _*M*_ for missense mutations and _*N*_ for nonsense mutations, and we excluded mutations classified as *‘*Silent*’, ‘*Intron*’, ‘*3*’* UTR*’, ‘*5*’* UTR*’, ‘*IGR*’, ‘*lincRNA*’*, and *‘*RNA*’*. For computational efficiency, we restricted our analysis to the 500 most frequently mutated genes across samples – a criterion that is typically used in other mutual exclusivity analyses – yielding a total of 124, 750 gene pairs that we analyze. We obtained background mutation rate distributions ℙ (*P*_*i*_) for each gene and mutation type (missense, nonsense) using CBaSE [V1.2] [75]. We emphasize that DIALECT could also be run with other methods for estimating background mutation rate distributions such as MutSigCV2 [40] or Dig [67].

#### Mutual exclusivity

DIALECT identified between 5 and 14 gene pairs in each of the five different cancer types. In contrast, DISCOVER, DISCOVER*, and Fisher*’*s exact test reported a higher number of pairs across all cancer subtypes, including over 300 pairs for colon adenocarcinoma and rectum adenocarcinoma (COADREAD) and uterine corpus endometrial carcinoma (UCEC). This pattern suggests that these methods may be prone to identifying interactions between genes with high numbers of mutations, many of which are likely passengers. Thus, for each method, we next evaluated the fraction of *“*suspicious*”* genes, or genes that are likely not driver genes as annotated by [40], in the mutually exclusive pairs identified by each method. Such suspicious genes have high numbers of passenger mutations, and are commonly identified or removed from the analyses by existing mutual exclusivity methods. We find that DIALECT does not identify pairs with suspicious genes, while 5-10% of the pairs identified by DISCOVER, DISCOVER*, and Fisher*’*s exact test contain suspicious genes (Figure 3A). As another assessment, we find that DIALECT identifies gene pairs with lower average mutation frequencies compared to gene pairs identified by DISCOVER, DISCOVER*, and Fisher*’*s exact test (Figure 3B). Genes with high mutation frequencies are often falsely identified by other methods, and contribute to the larger number of gene pairs identified by these methods. These analyses indicate that DIALECT does not identify mutual exclusivity between likely passenger genes with large numbers of mutations, in contrast DISCOVER, DISCOVER*, and Fisher*’*s exact test which often identify suspicious or highly mutated genes.

**Figure 3:**
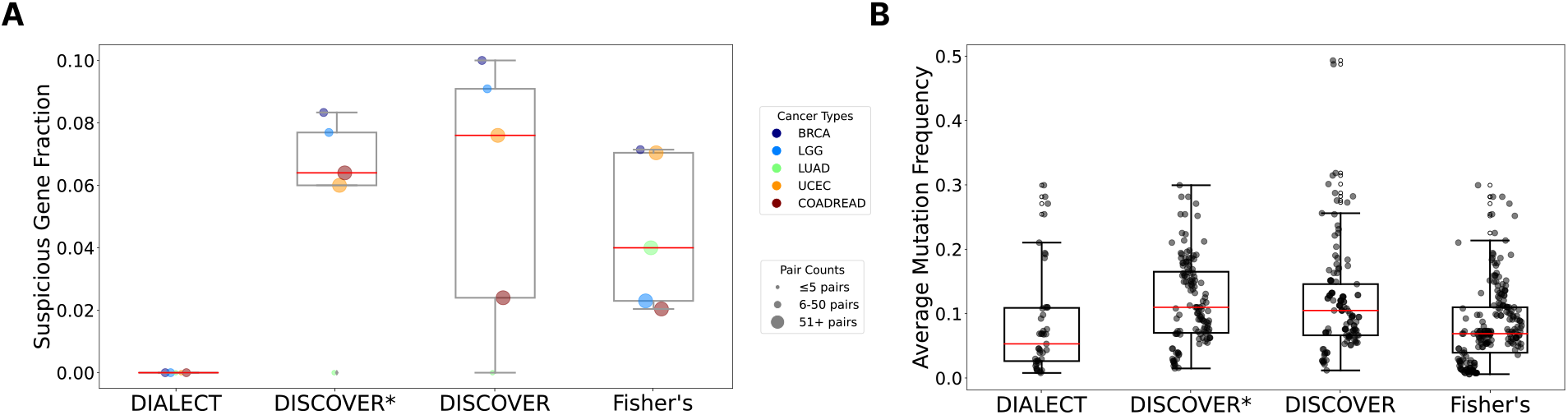
Comparison of pairs of genes identified by DIALECT, DISCOVER, and Fisher*’*s exact test for 5 cancer subtypes in The Cancer Genome Atlas (TCGA). **(A)** Suspicious gene fractions, or the fraction of gene pairs where at least one gene is in a list of *“*suspicious*”* genes that are likely not driver genes, as annotated in [40], for DIALECT, DISCOVER, DISCOVER*, and Fisher*’*s exact test. DISCOVER* is a variant of DISCOVER that is run separately on missense and nonsense mutations, similar to DIALECT. We select all gene pairs with *q*-value less than 0.05 for DISCOVER, DISCOVER*, and Fisher*’*s exact test. **(B)** The average mutation frequency of the two genes in each gene pair identified by DIALECT, DISCOVER, DISCOVER*, and Fisher*’*s exact test.

Focusing on breast cancer, the largest cohort in the dataset with *N* = 1084 patients, we observed (Table 1) that the gene pairs with the highest rates of mutual exclusivity, i.e. the pairs with largest log-odds estimated by DIALECT, are comprised of genes that are reported as drivers in breast cancer. Pairs such as CDH1_N:TP53_M (DIALECT *p*-value = 0.002) and AKT1_M:PIK3CA_M (DIALECT *p*-value = 0.015) have been found to reflect distinct functional modules within breast cancer, e.g. *TP53, CDH1, AKT1*, and *PIK3CA* are all known breast cancer driver genes [57, 37, 62].

**Table 1:** Mutually exclusive pairs of mutations identified by DIALECT, DISCOVER*, and Fisher*’*s Exact Test on TCGA breast cancer (BRCA) data. Higher LLR, lower q-values, and lower p-values indicate stronger mutual exclusivity. Suspicious genes are shown in bold. Pairs uniquely identified by a method are shown with ‡.

| DIALECT |  | DISCOVER* |  | Fisher’s Exact Test |  |
| --- | --- | --- | --- | --- | --- |
| Pair | LLR | Pair | q-value | Pair | p-value |
| CDH1_N:TP53_M | 14.728 | PIK3CA_M:TP53_M | $4.45 * 10^{-7}$ | CDH1_N:TP53_M | $7.46 * 10^{-4}$ |
| TP53_M:TP53_N | 12.132 | TP53_M:TP53_N | $9.57 * 10^{-6}$ | PIK3CA_M:TP53_M | $1.08 * 10^{-3}$ |
| PIK3CA_M:TP53_N | 11.153 | CDH1_N:TP53_M | $2.13 * 10^{-5}$ | TP53_M:TP53_N | $1.39 * 10^{-3}$ |
| AKT1_M:PIK3CA_M | 10.463 | PIK3CA_M:TP53_N | $4.98 * 10^{-5}$ | PIK3CA_M:TP53_N | $1.56 * 10^{-3}$ |
| PIK3CA_M:TP53_M | 9.933 | AKT1_M:PIK3CA_M | $4.44 * 10^{-4}$ | AKT1_M:PIK3CA_M | $1.84 * 10^{-3}$ |
| MAP3K1_N:TP53_M | 8.877 | MAP3K1_M:TP53_M | $3.54 * 10^{-3}$ | MAP3K1_N:TP53_M | $1.08 * 10^{-2}$ |
| NCOR1_N:TP53_M | 7.049 | MAP3K1_N:TP53_M | $5.24 * 10^{-3}$ | MAP3K1_M:TP53_M | $1.61 * 10^{-2}$ |
| ARID1A_N:TP53_M | 6.239 | FOXA1_M:TP53_M | $6.88 * 10^{-3}$ | FOXA1_M:TP53_M | $2.43 * 10^{-2}$ |
| FOXA1_M:TP53_M | 5.813 | AKT1_M:**TTN_M** | $1.01 * 10^{-2}$ | NCOR1_N:TP53_M | $2.82 * 10^{-2}$ |
| MYH9_M:TP53_M | 4.750 | MYH9_M:TP53_M | $1.92 * 10^{-2}$ | CBFB_M:TP53_M | $3.58 * 10^{-2}$ |
| MAP3K1_M:TP53_M | 4.728 | NCOR1_N:TP53_M | $3.78 * 10^{-2}$ | MYH9_M:TP53_M | $3.66 * 10^{-2}$ |
| CBFB_M:TP53_M | 3.898 | AHNAK2_M:TP53_M<sup>‡</sup> | $4.44 * 10^{-2}$ | AKT1_M:**TTN_M** | $4.34 * 10^{-2}$ |
| STAB2_M:TP53_M<sup>‡</sup> | 3.676 | | | GREB1L_M:TP53_M<sup>‡</sup> | $4.55 * 10^{-2}$ |
| AKT1_M:TP53_N | 3.519 | | | ARID1A_N:TP53_M | $4.55 * 10^{-2}$ |

In contrast, DISCOVER* and Fisher*’*s Exact Test identify spurious pairs that contain at least one *“*suspicious*”* gene. In particular, both DISCOVER* and Fisher*’*s exact test identify the pair AKT1_M:TTN_M. *TTN* has many random passenger mutations due to its extraordinary length and likely does not contain any driver mutations [39, 40]. The identification of the suspicious gene *TTN* by Fisher*’*s exact test agrees with its low specificity as we demonstrated in simulations (Figure 2E).

DISCOVER and DISCOVER* are particularly prone to identifying interactions between genes with high mutation rates, an issue exacerbated in types like COADREAD and UCEC which exhibit higher background mutation rates. In particular, COADREAD and UCEC samples typically exhibit a higher number of mutated genes per sample (median of 78.5 genes per sample for COADREAD and 57.5 genes per sample for UCEC) [42]. DISCOVER and DISCOVER* report over 500 significant pairs in COADREAD and over 1000 pairs in UCEC. In contrast, DIALECT identifies a far more selective 8 and 5 mutually exclusive pairs for COADREAD (Table S2) and UCEC (Table S3), respectively.

DIALECT also identifies novel mutual exclusivity between driver mutations that were not identified by existing methods. In particular, DIALECT identifies mutual exclusivity between STAB2_M:TP53_M. This pair was not identified by DISCOVER* or Fisher*’*s exact test (Figure 4, Table 1) due to the low mutation rate of *STAB2. STAB2* overexpression has been observed to cause increased tumor metastasis rates [29] and poor tumor prognosis [79], and may explain the observed mutual exclusivity between missense mutations in *TP53* and *STAB2*. These examples demonstrate how by modeling driver and passenger mutations separately, DIALECT is able to identify novel driver mutations and mutual exclusivity relations that are missed by current approaches.

**Figure 4:**
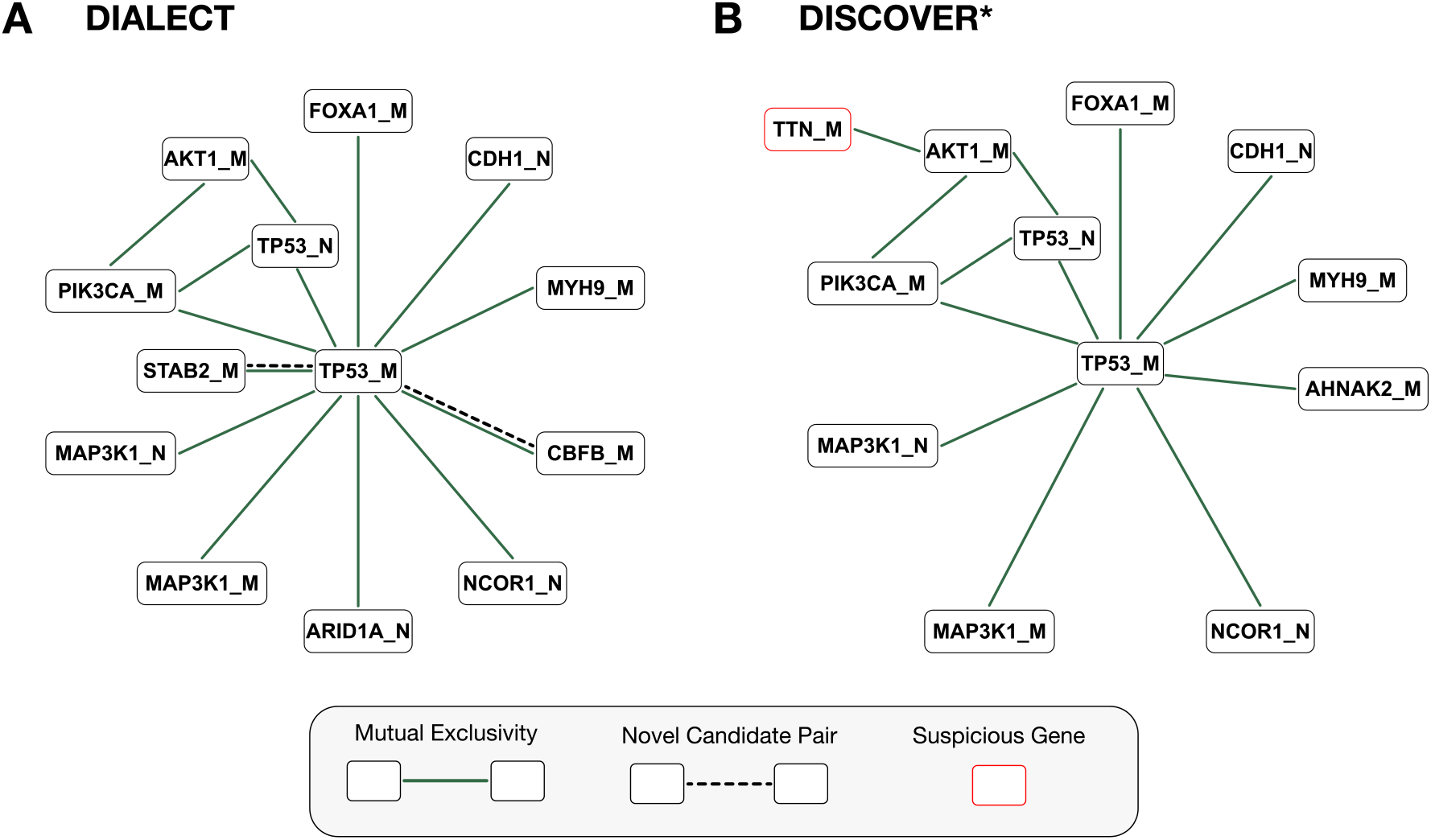
Mutually exclusive pairs of genes detected by DIALECT and DISCOVER* in breast cancer (BRCA). **(A)** Network of mutually exclusive gene pairs identified by DIALECT, where nodes represent genes, solid edges indicate mutual exclusivity between driver mutations, and dashed edges indicate novel gene pairs not identified in prior literature. **(B)** Network of mutually exclusive gene pairs identified by DISCOVER*. Red highlighted node indicates *“*suspicious*”* gene as annotated by [40].

## 4 Discussion

We introduce DIALECT, a method for identifying dependencies between pairs of *driver* mutations from somatic mutations counts. DIALECT explicitly models the observed somatic mutation counts as a sum of driver mutations and passenger mutations, in contrast to nearly all other methods which conflate drivers with passengers in a gene by *binarize* the mutation events in a gene. DIALECT models the distribution of driver mutations using a latent variable model while accounting for passenger mutations by incorporating existing background mutation rate (BMR) models. We derive an expectation maximization (EM) algorithm to estimate the parameters of our model which describe the degree of mutual exclusivity or co-occurrence between driver mutations. We demonstrate that DIALECT has improved performance compared to the standard mutual exclusivity and co-occurrence tests on simulated and real data.

Our approach for jointly modeling passenger and driver mutations can be readily extended in several directions. First, there are many methods for modeling BMRs, with each method having different strengths and weaknesses. In large-scale cancer studies, a standard practice is to form a *“*consensus*”* list of driver genes using BMRs estimated by different methods. Likewise, we imagine that it would be beneficial to run DIALECT with different BMR models in order to form a consensus list of mutually exclusive driver mutations. Second, although DIALECT allows for sample-specific BMRs (as demonstrated in simulations), existing tools do not readily output sample-specific BMRs for real data. Thus it would be useful to evaluate DIALECT using accurate sample-specific BMRs on a large-scale cohort. Similarly, DIALECT assumes that each tumor sample has an equal probability of a driver mutation, and we show in simulations that DIALECT has large power even when this assumption does not hold (i.e. when there is *model mis-specification*). Nevertheless, it may be useful to derive a more general model that incorporates sample-specific driver probabilities. Third, in the present work we used DIALECT to identify mutual exclusivity between driver mutations in real data, which provides a signal that the driver mutations perturb different biological pathways. Preliminary analysis suggests that there is no statistically significant co-occurrence in the TCGA data consistent with previous studies [10], but further analysis of this issue is necessary. Finally, we believe that our novel approach for separately modeling driver and passenger mutations would be advantageous for other problems in cancer genomics, particularly for learning cancer progression models (CPMs) which describe patterns in driver mutation accumulation over time [46, 64, 19,1,11, 54,66,47, 33].

## Availability

DIALECT is available online at https://github.com/raphael-group/dialect.

## 5 Acknowledgments

This research is supported by NIH/NCI grants U24CA248453 and U24CA264027 to B.J.R. U.C. was supported by NSF GRFP DGE 2039656 and the Siebel Scholars program. We thank Donate Weghorn for modifying CBaSE to output sample-specific background mutation distributions, and we thank Madelyne Xiao for work on a previous iteration of the model.

## Footnotes

1 One notable exception are tumor suppressor genes where both copies of the gene are typically inactivated (*“*two hit hypothesis*”*). However, it is common for one of these mutations to be a copy number aberration.

