## supplementary information for "A latent variable model for evaluating mutual exclusivity and co-occurrence between driver mutations in cancer"

### A Estimating driver mutation rates with DIALECT

We evaluated the driver mutation rates learned by DIALECT in order to demonstrate that DIALECT learns realistic parameters. Specifically, we used DIALECT to learn the driver mutation probability $\pi$ for different genes and mutation events across 1,008 breast cancer (BRCA) samples. DIALECT identifies 39 genes with FDR corrected $p$-value less than 0.05. We observed (Table S1) that the genes with largest driver mutation probabilities $\pi$, as estimated by DIALECT, include many genes that have been characterized as breast cancer driver genes, such as *TP53* ($q < 0.001$) [61] and *PIK3CA* ($q < 0.001$) [50]. Importantly, DIALECT does not estimate large driver mutation probabilities $\pi$ for the so-called ”suspicious” genes identified by [40], which is a list of genes that have a large number of somatic mutations but are likely not important for cancer development, e.g. the long genes *TTN* and *MUC16*. We emphasize that existing methods for driver gene identification, e.g. CBaSE [75, 40, 67] do not estimate the probability $\pi$ that a gene has a driver mutation in a given sample, in contrast to DIALECT.

| BRCA | | LAML | | LGG | |
|---|---|---|---|---|---|
| Gene | $\pi$ | Gene | $\pi$ | Gene | $\pi$ |
| PIK3CA_M | 0.315 | DNMT3A_M | 0.208 | IDH1_M | 0.776 |
| TP53_M | 0.206 | FLT3_M | 0.106 | TP53_M | 0.410 |
| TP53_N | 0.053 | IDH2_M | 0.106 | ATRX_N | 0.125 |
| KMT2C_N | 0.036 | IDH1_M | 0.096 | CIC_M | 0.106 |
| CDH1_N | 0.034 | NRAS_M | 0.081 | PIK3CA_M | 0.070 |
| AKT1_M | 0.025 | TET2_N | 0.075 | EGFR_M | 0.065 |
| VPS13C_M | 0.022 | DNMT3A_N | 0.056 | TP53_N | 0.048 |
| FOXA1_M | 0.022 | TP53_M | 0.053 | ATRX_M | 0.040 |
| ERBB2_M | 0.022 | PTPN11_M | 0.050 | IDH2_M | 0.038 |
| MAP3K1_N | 0.021 | KRAS_M | 0.049 | NOTCH1_M | 0.033 |

**Table S1:** Top 10 Genes and their Pi Values for Breast Invasive Carcinoma (BRCA), Acute Myeloid Leukemia (LAML), and Lower Grade Glioma (LGG). The suffixes ‘_M‘ and ‘_N‘ denote missense and nonsense mutations, respectively.

### B DIALECT mutual exclusivity result tables

| Pair | LLR |
|---|---|
| KRAS_M:BRAF_M | 42.904 |
| TP53_M:TP53_N | 26.657 |
| APC_N:BRAF_M | 25.290 |
| TP53_M:BRAF_M | 13.850 |
| TP53_M:PIK3CA_M | 13.130 |
| KRAS_M:COL7A1_M | 10.686 |
| TP53_M:CHD8_M | 10.545 |
| TP53_M:DSPP_M | 10.155 |

**Table S2:** Mutually exclusive gene pairs identified by DIALECT in colon and recum adenocarcinoma (COADREAD).

| Pair | LLR |
|---|---|
| TP53_M:CTNNB1_M | 29.599 |
| TP53_M:KRAS_M | 20.475 |
| TP53_M:ARID1A_N | 19.767 |
| PTEN_M:TP53_M | 18.349 |
| TP53_M:PTEN_N | 12.914 |

**Table S3:** Mutually exclusive gene pairs identified by DIALECT in uterine corpus endometrial carcinoma (UCEC).

| Pair | LLR |
|---|---|
| IDH1_M:EGFR_M | 101.862 |
| IDH1_M:IDH2_M | 61.625 |
| TP53_M:CIC_M | 41.478 |
| IDH1_M:PTEN_M | 29.242 |
| TP53_M:EGFR_M | 25.052 |
| CIC_M:ATRX_N | 16.733 |
| IDH1_M:MYOCD_M | 13.221 |
| ATRX_N:EGFR_M | 12.431 |
| IDH1_M:DHX30_M | 10.865 |
| IDH1_M:FLG_M | 10.797 |
| TP53_M:IDH2_M | 10.252 |
| IDH1_M:ABLIM3_M | 10.217 |

**Table S4:** Mutually exclusive gene pairs identified by DIALECT in brain lower grade glioma (LGG).

| Pair | LLR |
|---|---|
| TP53_M:TP53_N | 31.518 |
| KRAS_M:EGFR_M | 21.707 |
| KRAS_M:BRAF_M | 20.458 |
| TP53_M:KRAS_M | 13.123 |

**Table S5:** Mutually exclusive gene pairs identified by DIALECT in lung adenocarcinoma (LUAD).

### C EM implementation details

We use a total of 8 initializations for the $\tau_{00}$, $\tau_{01}$, $\tau_{10}$, $\tau_{11}$ when running EM: five random, one each for co-occurrence, mutual exclusivity, and independence scenarios. Each initialization followed specific formulas to set $\tau_{00}, \tau_{01}, \tau_{10}, \tau_{11}$.

**Random Initializations.** Five random initializations were created by selecting linearly spaced values for $\tau_{00}$ between 0.8 and 1, given that most gene pairs generally did not have more than 10% of cases where either gene had at least one mutation.

**Co-occurrence Initialization.** This initialization represented cases where both genes in a pair had a high tendency to mutate simultaneously. It was defined as:

$$\begin{aligned}
\tau_{00} &= 1 - \max(\widehat{\pi}, \widehat{\pi}'), \\
\tau_{10} &= 0 \text{ if } \widehat{\pi} < \widehat{\pi}' \text{ else } \widehat{\pi} - \widehat{\pi}', \\
\tau_{01} &= 0 \text{ if } \widehat{\pi}' < \widehat{\pi} \text{ else } \widehat{\pi}' - \widehat{\pi}, \\
\tau_{11} &= \min(\widehat{\pi}, \widehat{\pi}').
\end{aligned}$$

**Mutual Exclusivity Initialization.** This initialization represented cases where the mutations in the genes were mutually exclusive. It was defined as:

$$\begin{aligned}
\tau_{00} &= 1 - (\widehat{\pi} + \widehat{\pi}'), \\
\tau_{10} &= \widehat{\pi}, \\
\tau_{01} &= \widehat{\pi}', \\
\tau_{11} &= 0.
\end{aligned}$$

**Independence Initialization.** This initialization corresponded to the scenario where the mutations in the genes occurred independently of each other. It was defined as:

$$\begin{aligned}
\tau_{00} &= (1 - \widehat{\pi}) \cdot (1 - \widehat{\pi}'), \\
\tau_{10} &= \widehat{\pi} \cdot (1 - \widehat{\pi}'), \\
\tau_{01} &= (1 - \widehat{\pi}) \cdot \widehat{\pi}', \\
\tau_{11} &= \widehat{\pi} \cdot \widehat{\pi}'.
\end{aligned}$$
